# Digital Kennison: A bioinformatics pipeline for rapid mapping of sequences to the *Drosophila melanogaster* Y chromosome

**DOI:** 10.64898/2026.07.15.738766

**Authors:** Fabiana Uno, A. Bernardo Carvalho

## Abstract

The *Drosophila melanogaster* Y chromosome is currently known to contain 13 single-copy protein-coding genes, six of which are essential for male fertility, as well as several non-coding genes and abundant repetitive DNA. Localization of Y-linked sequences has traditionally relied on labor-intensive crosses using Kennison’s translocation strains, which map Y-linked loci by generating flies deficient for each of the six Y-chromosome fertility regions (ks-1, ks-2, kl-1, kl-2, kl-3, and kl-5). Here we present Digital Kennison, a computational pipeline that recasts this classical mapping strategy as a sequence-based analysis. The pipeline queries eight genomic databases derived from Kennison’s strains using BLAST and read coverage, assigning sequences to fertility regions with a calibrated confidence score. We benchmarked the method on 60 Y-linked sequences spanning all six regions, including single-copy protein-coding genes, *Mst77Y* family members, non-coding RNAs, and the centromere. Digital Kennison achieved 97% precision while resolving challenging cases, including boundary-spanning genes (*PRY* and *Ppr-Y*), fragmented *Mst77Y* copies, and *FDY*, which has a closely related autosomal paralog. Beyond validating known localizations, the pipeline localized the unmapped gene *CG41561* to the kl-1region and reassigned the transcript *CR40629-RC* from the kl-2 region to kl-5. It also localized 7 of 16 recently transferred Y-linked sequences described by Tobler et al. (2017), including 4 with high confidence. Applied to 904 small R6 scaffolds, Digital Kennison assigned 75% to fertility regions, including five currently annotated as autosomal-pericentromeric. Digital Kennison reduces sequence localization from weeks of genetic crosses to minutes of computation while preserving the power of classical translocation mapping.

**Article summary:** The *Drosophila melanogaster* Y chromosome is difficult to study because it consists largely of repetitive, non-recombining DNA. Researchers have traditionally mapped Y-linked genes using slow, labor-intensive genetic crosses. Here we introduce Digital Kennison, a computational pipeline that replicates this classical mapping strategy using DNA sequence data instead of live flies. By comparing a query sequence against genomic databases built from fly strains, the pipeline assigns it to one of six Y-chromosome regions and reports a confidence score. Tested on 60 known sequences, it achieved 97% precision, corrected an annotation error, and mapped previously unplaced sequences in minutes rather than weeks.

## Introduction

The *Drosophila melanogaster* Y chromosome is essential for male fertility but plays no role in sex determination (Bridges, 1916). It is entirely heterochromatic and spans approximately 40 Mb (Gatti & Pimpinelli, 1983), although less than 20 Mb has been assembled to date (Chang & Larracuente, 2019; Shukla et al., 2025). The Y is divided into a long arm (Y^L^) and a short arm (Y^S^) and, unlike the euchromatic chromosome arms, neither forms polytene chromosomes nor undergoes meiotic recombination. These features have historically limited the mapping of Y-linked genes.

Genome sequencing could, in principle, overcome these limitations, but the extreme abundance of repetitive DNA on the Y chromosome has led to fragmented, incomplete assemblies. In Release 6 (Hoskins et al., 2015; Öztürk-Çolak et al., 2024), the largest scaffold assigned to the Y chromosome (∼3.7 Mb) is not a single continuous sequence, but rather a collection of shorter contigs stitched together for computational convenience. Long-read sequencing substantially improved Y-chromosome assembly (Krsticevic et al., 2015, Carvalho et al., 2015, Chang & Larracuente, 2019), but progress remains severely limited by a strong sequencing bias against simple satellites such as (AAGAG)n, which are very abundant on the Y chromosome (Carvalho et al., 2026).

Given these challenges, it is hardly surprising that the Y remains the least well-characterized chromosome in the *D. melanogaster* genome. In addition to rDNA and a few repetitive gene families, only a limited number of protein-coding genes have been confidently assigned to the Y chromosome (Gepner & Hays, 1993; Carvalho et al., 2015 and references cited therein; Mahajan & Bachtrog, 2017). Its complete gene content, structural organization, and the precise localization of many Y-linked sequences remain largely unresolved.

Perhaps due to its difficulty and enigmatic nature, the Y chromosome has attracted the attention of several prominent geneticists. Bridges (1916) showed that the Y is not involved in sex determination but is essential for male fertility: X0 males are phenotypically normal but sterile. Stern (1929) later demonstrated that both arms (Y^L^ and Y^S^) are required for male fertility, establishing that the Y harbors multiple fertility factors distributed along its length.

These foundational insights were followed by decades of genetic research aimed at identifying and mapping the Y-linked fertility factors. Brosseau (1960) posed two central questions: “*how many genes are there in each complex and can they be linearly ordered?*“ While linear ordering proved feasible, determining the exact number of genes has been more challenging. Brosseau (1960) originally identified at least two fertility genes on the short arm (*ks-1* and *ks-2*) and proposed five on the long arm (*kl-1* to *kl-5*). A key difficulty in these studies is that the Y chromosome does not recombine, precluding classical genetic mapping. Geneticists were therefore forced to rely on X-ray-induced deletions and complementation analysis (Brosseau, 1960; Hazelrigg et al., 1982). A major limitation of this strategy is the instability of deletion stocks: maintaining male-sterile Y chromosomes requires a helper Y chromosome (e.g., an attached XY), rendering the mutated Y dispensable and prone to accumulating additional rearrangements over time. Indeed, Hazelrigg et al. (1982) concluded that the original Brosseau lines could no longer be reliably used as tester chromosomes.

A breakthrough came with Kennison (1981), who generated both fertile and sterile X–Y translocations with one breakpoint in the Y chromosome and the other in a nonessential region of the X heterochromatin. Analysis of the sterile translocations refined the functional map of the Y chromosome, confirming two fertility factors on the short arm and four on the long arm (kl-4 was not supported as a separate locus; the same conclusion was independently reached by Hazelrigg et al. (1982)). The fertile translocations proved even more valuable. By crossing lines carrying different Y-chromosome breakpoints, defined Y deficiencies could be generated in F1 males at will. Unlike deletion stocks, the parental translocation lines were maintained under fertility selection and therefore remained stable over time. Indeed, Carvalho et al. (2000) repeated Kennison’s original crosses nearly two decades later and obtained the same results. This system effectively solved the instability problem of reference strains that had limited earlier deletion-based approaches.

Kennison’s work benefited from the then-unpublished cytogenetic method of Gatti & Pimpinelli (1983), who developed specialized mitotic chromosome preparations that subdivided the Y chromosome into 25 heterochromatic bands (h1–h25), creating a map that remains in use to this day. During the same period, studies using Kennison’s lines demonstrated that the Y chromosome fertility factors are conventional protein-coding genes (Hardy et al., 1981; Goldstein et al., 1982) a finding in which Dan Lindsley, who supervised Kennison’s doctoral work, played a central role. Although controversial at the time (Hennig, 1993), this result set the stage for the molecular identification of Y-linked genes.

Kennison translocation lines have become the standard tool for mapping Y-linked sequences (e.g., Livak, 1984; Gepner & Hays, 1993; Danilevskaya et al., 1993; Zhang & Stankiewicz, 1998; Carvalho et al., 2000, 2001, 2015). Using them, a Y-linked gene can be mapped to one of seven contiguous regions — corresponding to the six fertility factors plus the centromeric region between kl-1 and ks-1 — whose boundaries correspond to translocation breakpoints mapped onto the heterochromatic bands h1-h25s (Figure 2). These regions are named after the fertility genes they encompass but may contain additional loci; for example, the kl-5 region contains the complete *kl-5* and *Pp1-Y1* genes and part of *PRY* (Carvalho et al., 2000, 2001).

**Figure 1.**
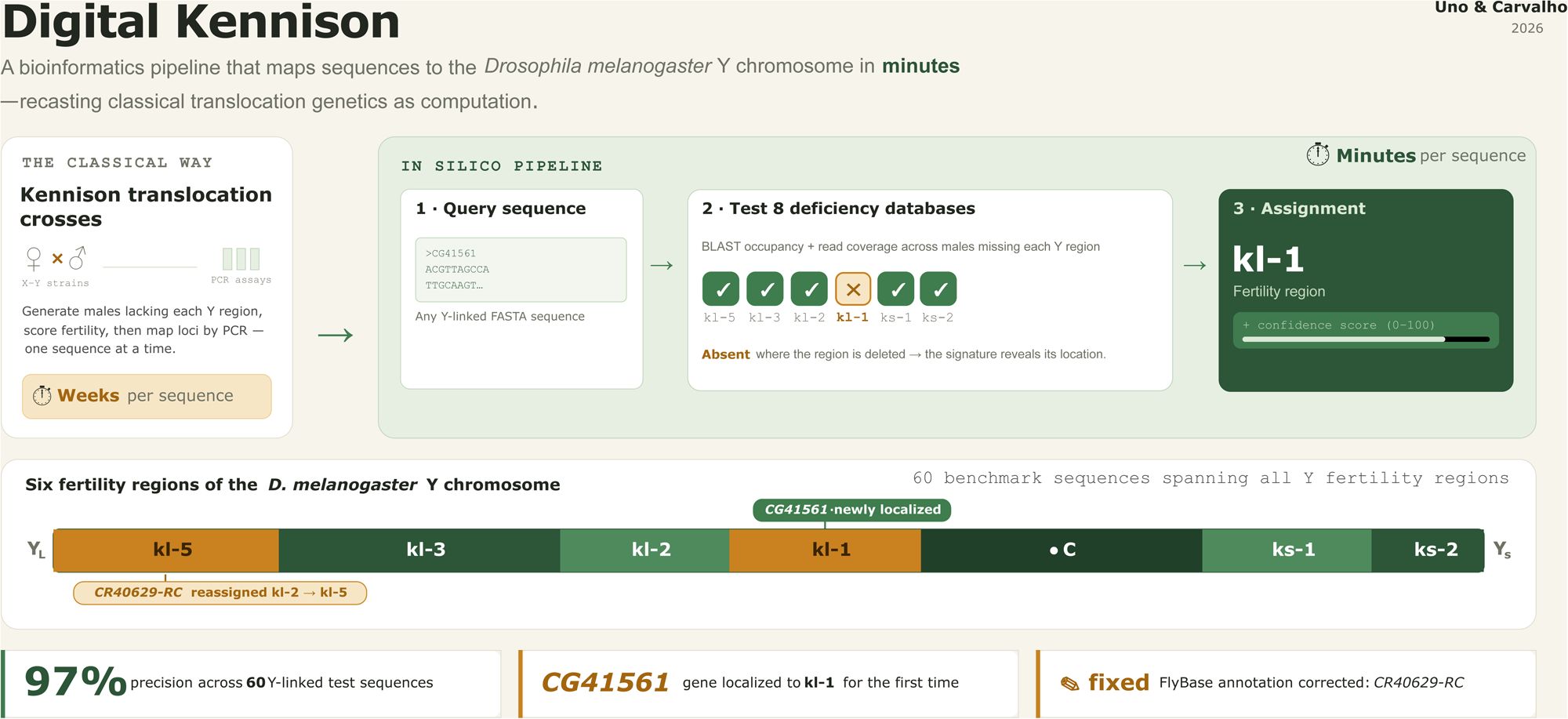
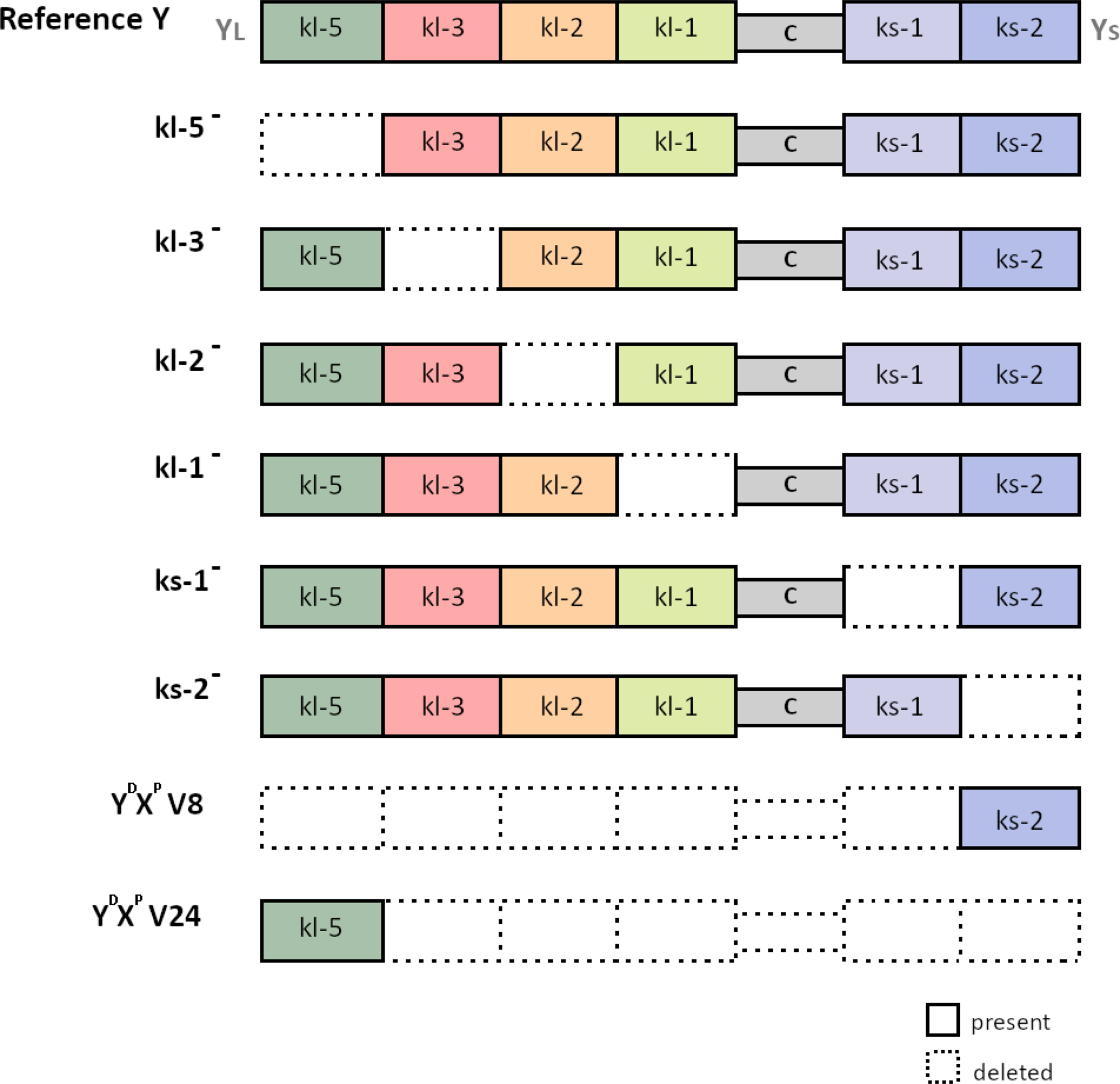
Y-chromosome deficiencies generated with the Kennison lines and employed by the Digital Kennison pipeline. The *D. melanogaster* Y chromosome (top) is partitioned into seven regions: kl-5, kl-3, kl-2, kl-1, ks-1, ks-2, and the centromere. Each genotype (kl-5*⁻,* kl-3*⁻*, etc.) lacks a defined Y-chromosome region (dashed outline), whereas Y^D^X^P^ V8 and Y^D^X^P^ V24 retain only the ks-2 and kl-5 regions, respectively. Comparisons between user-provided sequences and genomic databases derived from these lines allow Digital Kennison to infer the chromosomal location of Y-linked sequences. **ALT TEXT:** Schematic of the *Drosophila melanogaster* Y chromosome and the eight Y-deficiency genotypes. The reference chromosome is divided into seven fertility regions. Each deficiency genotype lacks one region, indicated by a dotted outline, while V8 and V24 retain only ks-2 or kl-5, respectively.

**Figure 2.**
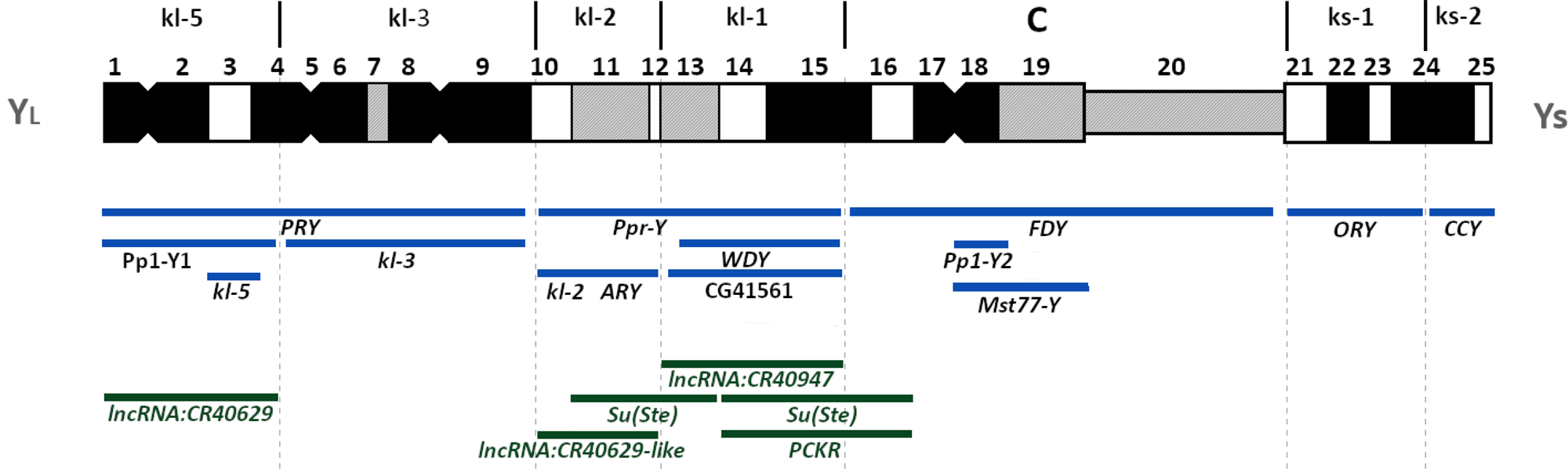
Organization of the *D. melanogaster* Y chromosome. The Y chromosome is subdivided into 25 heterochromatic bands (h1–h25), with the centromere region marked as “C”. Fertility factor regions are denoted as kl-1 through kl-5 on the long arm (Y^L^) and *ks-1* and *ks-2* on the short arm (Y^S^). The upper panel shows the cytological positions of well-characterized protein-coding Y-linked genes (blue), the highly repetitive *Su(Ste)* and *PCKR* arrays (green), and the non-coding transcripts *lncRNA: CR40947*, *lncRNA: CR40629*, and the *CR40629*-like paralog cluster (green). A full list of known Y-linked genes with cytological positions and references is provided in *Supplementary Table S6*. **ALT TEXT**: Linear map of the *Drosophila melanogaster* Y chromosome showing its 25 cytological bands, centromere, and seven fertility regions (kl-5 to ks-2). Blue bars indicate the positions of protein-coding Y-linked genes, including boundary-spanning genes, and green bars indicate the *Su(Ste)* and *PCKR* repeat arrays, three annotated non-coding RNAs, and the *CR40629*-like paralog cluster.

Since the advent of PCR, mapping with the Kennison lines has typically been performed by amplifying a target sequence against a panel of DNA templates, each derived from Y-deficient males lacking a specific Y region: a PCR-negative result indicates that the gene maps to the deleted region. While straightforward, this procedure requires obtaining several *Drosophila* strains from the Bloomington Stock Center, performing crosses, extracting DNA, and running PCR with appropriate controls — a process that can take weeks. Probably for this reason, several Y-linked genes and sequences have remained unmapped (e.g., *CG41561,* Mahajan & Bachtrog, 2017; and the Y-linked pseudogenes identified by Tobler et al., 2017).

Here we describe Digital Kennison, a bioinformatics pipeline that reframes Kennisońs classical mapping approach as a purely sequence computational analysis. The pipeline uses a fixed set of precomputed genomic databases derived from Y-deficient males that lack specific Y-chromosome regions to localize user-provided sequences. Although the underlying logic is straightforward, repetitive sequences, closely related autosomal paralogs, and genes spanning multiple fertility regions may complicate this inference. Digital Kennison addresses these challenges by integrating BLAST-based and read-based evidence to assign sequences to specific Y-chromosome regions. By converting Kennison’s mapping framework into a computational query, Digital Kennison enables fast mapping of newly identified sequences and genes, as well as cross-validation (and eventual correction) of existing annotations.

## Methods

### Generation of Y-region deficiency genotypes and DNA preparation

The DNA samples used for sequencing were the same as those used by Carvalho et al. (2015) to map the *FDY* gene by PCR. F1 males lacking defined regions of the Y chromosome were generated by Carvalho et al. (2000) using fertile X–Y translocation strains (Kennison lines), following the crossing scheme originally described by Kennison (1981). Details of the strains, crossing schemes, and resulting genotypes are listed in *Supplementary Table S1* and *Supplementary Table S2 in Supplementary File S1*. Genomic DNA was extracted by Carvalho et al. (2015) and stored at −20 °C until sequencing. No additional crosses or DNA extractions were performed for the construction of the deficiency databases. The Illumina sequencing data generated from these DNA samples have been deposited in the NCBI Sequence Read Archive under BioProject PRJNA1493028 (accessions: SRR39563621-SRR39563614). DNA samples generated specifically for the YGS analyses are described separately below.

### Genome sequencing and database construction

Samples were sequenced at Macrogen (Korea) in September 2024 using either the Illumina TruSeq Nano DNA protocol (for samples kl-5⁻, kl-3⁻, kl-1⁻, ks-1⁻, ks-2⁻, V8, and V24) or the Nextera DNA XT protocol (for sample kl-2⁻). Sequencing was performed in 151 bp paired-end mode with a target insert size of 350 bp, yielding a mean depth of ∼62× per sample (range 55–68×; *Supplementary Table S3 in Supplementary File S1*). Read quality was assessed with FastQC v0.11.9. Reads were processed with TrimGalore v0.6.6 and assembled de novo with SPAdes v3.15.5 (Bankevich et al., 2012) using default parameters. Contaminant sequences were removed with FCS-GX (Astashyn et al., 2024). Assembly completeness was assessed with BUSCO v5.8.0 against the Diptera_odb10 database (per-strain metrics are reported in *Supplementary Table S3 in Supplementary File S1*). The eight Kennison deficiency-panel BLAST databases used by the pipeline were constructed with makeblastdb (BLAST 2.12.0+). Full command lines and parameter values for all analyses described above are available in the project GitHub repository (see Data Availability). BLAST databases are restricted to contigs identified as Y-linked by the YGS method (Carvalho & Clark, 2013), suppressing alignments to autosomal paralogs. Details of the YGS analysis and database construction are described in *Supplementary Methods* (sections *S3.1 and S3.2*) *in Supplementary File S1*.

Read databases for the read-based classification module were prepared from the same trimmed paired-end datasets, with autosomal and X-linked reads removed to suppress signal from non-Y paralogs. Reads were aligned to the R6 reference (FlyBase release 6.64) using minimap2 (`-ax sr`); reads mapping to the major assembled chromosome arms (2L, 2R, 3L, 3R, 4, X) were excluded (both reads in each pair were removed). The remaining reads (mapping to Y, unplaced contigs, or unmapped) were used as input to the read-based classification module (*Supplementary Table S4 in Supplementary File S1*).

### The Digital Kennison Algorithm

The Digital Kennison pipeline (v36j; Zenodo: 10.5281/zenodo.21271417) takes as input a FASTA file containing one or more query sequences and returns, for each sequence, an assignment to a specific region of the *D. melanogaster* Y chromosome together with a confidence score. Classification is based on comparisons against a panel of eight genomic databases (the deficiency panel), each derived from males lacking defined Y-chromosome regions (Figure 1; *Supplementary Table S1 and Supplementary Table S2 in Supplementary File S1*). Digital Kennison integrates two independent lines of evidence. The first is BLAST-based occupancy (the percentage of non-N query positions covered by alignments in each deficiency database), which measures the presence or absence of the query sequence across assembled contigs from the deficiency panel. The second is read-based coverage, obtained by mapping the unassembled sequencing reads from the same strains directly to the query. For optimal performance, input sequences should first be screened for Y-linkage using the YGS method (Carvalho & Clark, 2013), the chromosome quotient (CQ) method (Hall et al., 2013), or a comparable approach. Non-Y sequences are not assigned to fertility regions and are returned as *Undetermined*.

#### BLAST-based classification

For each query sequence, the pipeline searches each deficiency database using BLASTn and computes occupancy and the total number of hits. Because a single identity threshold may either exclude genuine matches or admit paralogous hits, each query is evaluated across a range of thresholds (95–100% by default). The optimal threshold is selected by maximizing a weighted occupancy score that penalizes excessive numbers of hits (hit-weight factor). Sequences exhibiting anomalous occupancy patterns, such as uniformly low signal or autosomal contamination, are flagged during pre-filtering and excluded from further analysis (*Supplementary Methods, section S3.3.2 in Supplementary File S1*). The remaining occupancy profiles are compared with the expected presence/absence patterns of each fertility region (Table 1). Queries displaying clear binary signatures (presence ≥80% and absence ≤10%) are classified directly. In contrast, more complex cases are handled through specialized procedures, including detection of boundary-spanning genes, centromere-specific rules, tiered outlier analysis, and assembly-aware correction of fragmentation artifacts (*Supplementary Methods, section S3.3 in Supplementary File S1*).

**Table 1.** Expected presence/absence signatures of Y-chromosome fertility regions across the deficiency panel. For each fertility region (rows), the matrix gives the expected presence (1) or absence (0) of a sequence located in that region across the eight strain databases (columns). The deficiency genotypes kl-5⁻ through ks-2⁻ each lack a single fertility region, while V8 and V24 retain only the ks-2 and kl-5 regions, respectively. The pipeline assigns each query sequence to a region by matching its observed occupancy pattern across the panel to one of these expected signatures.

| region | kl-5minus | kl-3minus | kl-2minus | kl-1minus | ks-1minus | ks-2minus | V8 | V24 |
| --- | --- | --- | --- | --- | --- | --- | --- | --- |
| kl-5 | 0 | 1 | 1 | 1 | 1 | 1 | 0 | 1 |
| kl-3 | 1 | 0 | 1 | 1 | 1 | 1 | 0 | 0 |
| kl-2 | 1 | 1 | 0 | 1 | 1 | 1 | 0 | 0 |
| kl-1 | 1 | 1 | 1 | 0 | 1 | 1 | 0 | 0 |
| ks-1 | 1 | 1 | 1 | 1 | 0 | 1 | 0 | 0 |
| ks-2 | 1 | 1 | 1 | 1 | 1 | 0 | 1 | 0 |
| Centromere | 1 | 1 | 1 | 1 | 1 | 1 | 0 | 0 |

#### Read-based classification and integration

BLAST-based classification depends on the quality of the genome assemblies used to construct the deficiency databases and may fail when repetitive sequences are fragmented, misassembled, or absent. To provide an assembly-independent source of evidence, Digital Kennison also performs a read-based analysis using the unassembled Illumina data from each Y-deficient male. Paired-end Illumina reads are mapped to the query sequences with minimap2 (Li, 2018) using the prefiltered read datasets described above (*Supplementary Table S4, in Supplementary File S1*). Coverage breadth (the proportion of the query sequence supported by mapped reads) is then used to infer presence or absence in each deficiency background, following the same conceptual framework as the BLAST-based occupancy analysis but applied directly to sequencing reads. BLAST and read-based classifications are generated in parallel and subsequently integrated into a unified classification. Concordant assignments increase the final confidence score, informative read-based classifications can resolve queries left undetermined by BLAST, and discordant results are either resolved in favor of the higher-confidence evidence or flagged for manual inspection. This dual-evidence framework is particularly valuable for sequences affected by assembly fragmentation or paralogous interference, for which BLAST-based classification alone may be unreliable (see Results: *FDY*, *Mst77Y*).

#### Confidence scoring

Each classification is assigned a confidence score (0–100) that summarizes the strength of the supporting evidence. The score is a weighted combination of four positive components — pattern sharpness (40%), genomic context (25%), identity threshold (20%), and hit consistency (15%) — and a fifth penalty term (classification-method quality) that downgrades scores derived from relaxed or fallback classification paths. The final score is further adjusted for classification difficulty and agreement with read-based evidence. Full details of the scoring scheme and parameter thresholds are provided in *Supplementary Methods, section S3.3, in Supplementary File S1*.

#### Code revision and debugging assistance

Portions of the Digital Kennison pipeline code were revised and debugged using Claude (Anthropic, Claude Opus 4.7) via iterative conversational prompting to identify bugs, refactor functions, and improve code clarity. All AI-suggested changes were reviewed, tested, and validated by the authors, who take full responsibility for the pipeline’s correctness and performance.

### Benchmark datasets and evaluation

To evaluate robustness across different sequence types, we performed systematic testing using three curated datasets: (i) 13 well-characterized single-copy protein-coding genes distributed across all six male fertility regions; (ii) 18 multi-copy protein-coding genes from the tandem *Mst77Y* array; and (iii) 29 Y-linked non-coding RNAs (*Supplementary Table S6 in Supplementary File S1*). Each dataset was analyzed under multiple parameter configurations to assess the effect of key pipeline components on classification accuracy (*Supplementary Table S5 in Supplementary File S1*). To evaluate the reliability of the confidence scores, each classification was labeled as Correct, Undetermined, or Incorrect based on agreement with the known chromosomal location, and precision and overall accuracy were calculated at successive confidence thresholds (*Supplementary Table S16 in Supplementary File S1*).

### Classification of unlocalized scaffolds

We analyzed unmapped scaffolds from the *D. melanogaster* Release 6 (R6) assembly using the Digital Kennison pipeline under optimized parameters (*Supplementary Methods, section S3.5.1 in Supplementary File S1*). Y-linkage assignments were independently validated by comparison with four *D. melanogaster* assemblies: Release 5 (R5), the heterochromatin-enriched PacBio assembly of Chang and Larracuente (2019), a de novo PacBio HiFi assembly generated from the LILAP reads of Jia et al. (2024) and a de novo Oxford Nanopore assembly generated from the reads of Kim et al. (2024). Scaffold correspondences were established by scaffold-to-contig alignment with minimap2 (`-ax asm5`). Y-linkage was confirmed with the YGS method (Carvalho & Clark, 2013), which scores each contig by the proportion of male-specific k-mers unmatched by female reads (P_VSC_UK) (*Supplementary Figure S3 in Supplementary File S1*). Autosomal- or X-linked scaffolds showing strong Y-linkage signal (P_VSC_UK ≥ 80%) in at least three of the four assemblies were flagged as candidates for reassignment to the Y chromosome (see *Supplementary Results, section S.4.4 in Supplementary File S1* for details).

We next applied Digital Kennison to two de novo genome assemblies (ONT and LILAP; *Supplementary Methods, section S3.5.1, in Supplementary File S1*). Unlike the highly curated Release 6 reference, these assemblies lack prior chromosomal or regional annotation of Y-linked scaffolds. Candidate Y-linked scaffolds were first identified using the YGS method (Carvalho & Clark, 2013), and repetitive sequences were masked with RepeatMasker v4.1.8 before pipeline classification.

### Classification of centromere contigs

To assess whether Digital Kennison can recover the expected centromeric locations, we analyzed the five *D. melanogaster* centromere contigs described by Chang et al. (2019), comprising the Y centromere and the four autosomal centromeres. These sequences were classified under a targeted comparison of hit-weighting settings (see Results and *Supplementary Results, section S4.2 in Supplementary File S1*).

### Application to recently transferred Y-linked sequences (Tobler et al. 2017)

Tobler et al. (2017) identified 21 incipient Y-linked sequence transfers in *D. melanogaster,* using the BDGP5 (R5) assembly. We merged four of these into a single consensus transfer corresponding to the *FDY* locus (a previously characterized ∼11 kb duplication; Carvalho et al., 2015) and excluded two sequences that failed the HSP chaining step, yielding 16 input sequences (*Supplementary Table S9 in Supplementary File S1*). To ensure compatibility with our R6-based pipeline, R5 coordinates were converted to Release 6 coordinates, and the resulting sequences were filtered for Y-linkage. Sequences were assigned to one of four categories — Resolved, Multicopy single region, Dispersed, or Insufficient evidence — based on the number of Y-linked hits, the dominant inferred location and its support, and the agreement between BLAST and read-based evidence (category definitions and aggregation thresholds in *Supplementary Methods, section S3.5.3 in Supplementary File S1*). The resulting file was used as input for the Digital Kennison pipeline.

### Figure preparation

Figures 1, 2, and S4 were prepared using Inkscape (v1.4.2; commit f4327f4, 2025-05-13).

## Results

### Pipeline performance and confidence calibration

We benchmarked the Digital Kennison pipeline on 60 Y-linked sequences, spanning single-copy genes, the Mst77Y array, and non-coding RNAs, distributed across all six fertility regions (Methods, Tables 2 and 3), and used this set to calibrate the confidence score for each assignment. Across the benchmark, 55 of 60 sequences (91.7%) were correctly classified, four were misclassified (6.7%), and one returned an ambiguous result (1.7%). At a confidence threshold of ≥55, the pipeline achieved 97.1% precision. Excluding *CR40629-RC* (a likely FlyBase annotation error discussed below), precision reached 100% for all thresholds ≥55 (*Supplementary Table S16 and Supplementary Figure S5 in Supplementary File S1*).For practical applications, we recommend interpreting scores ≥80 as suitable for unsupervised classification; scores between 55 and 80 as generally reliable but benefiting from independent confirmation; and scores <55 as requiring manual inspection or experimental confirmation, particularly for repetitive or multicopy sequences.

**Table 2.**
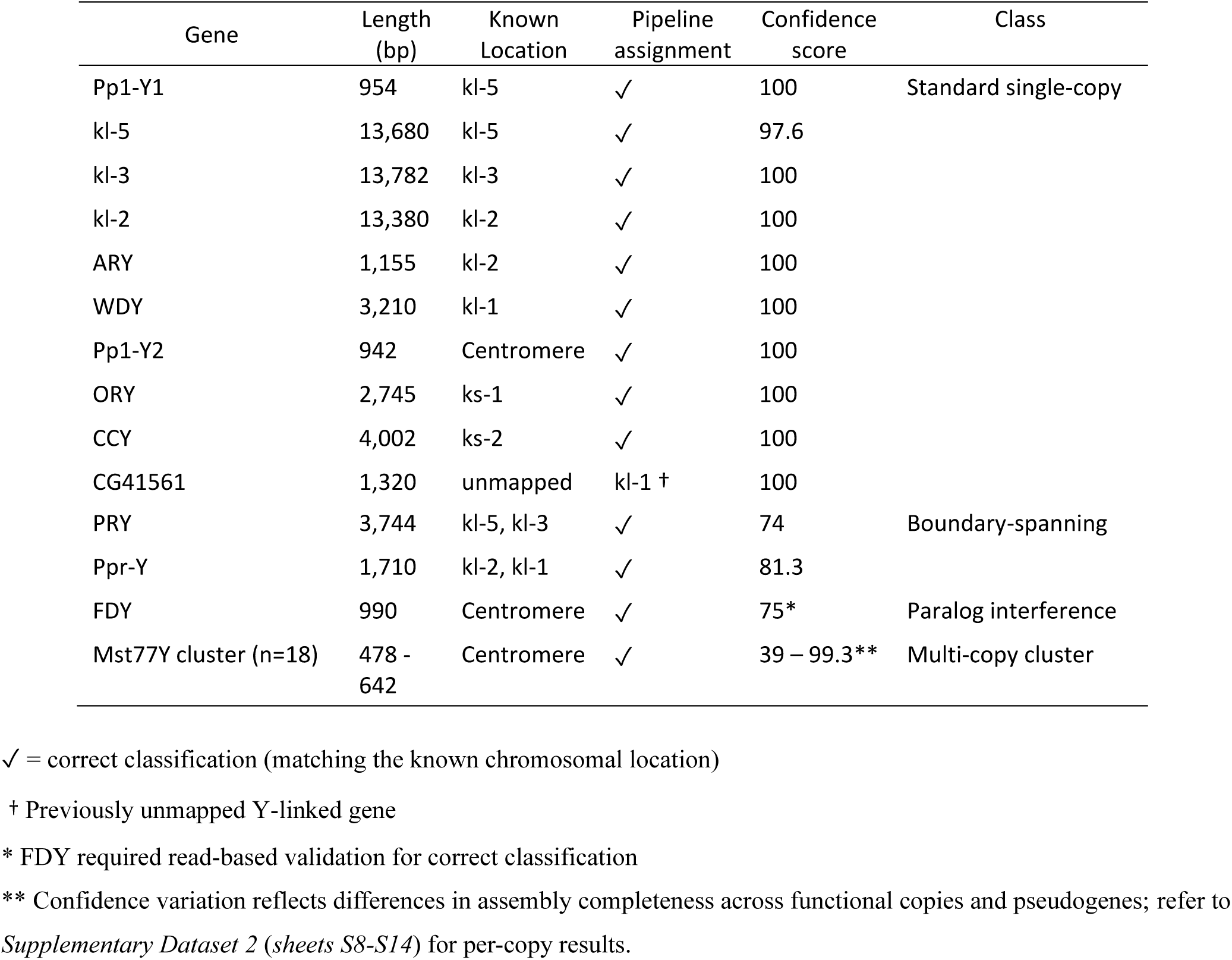
Classification performance of the Digital Kennison pipeline on known Y-linked protein-coding genes, grouped by sequence complexity class. For each gene, the table shows the sequence length, predicted chromosomal location assigned by the pipeline, and the known cytological location from the literature. The classification was run under optimized parameters. The confidence score reflects the strength of evidence supporting each classification, ranging from 0 to 100. For the *Mst77Y* gene cluster, the reported size range reflects multiple paralogs, and the confidence scores vary accordingly.

**Table 3.**
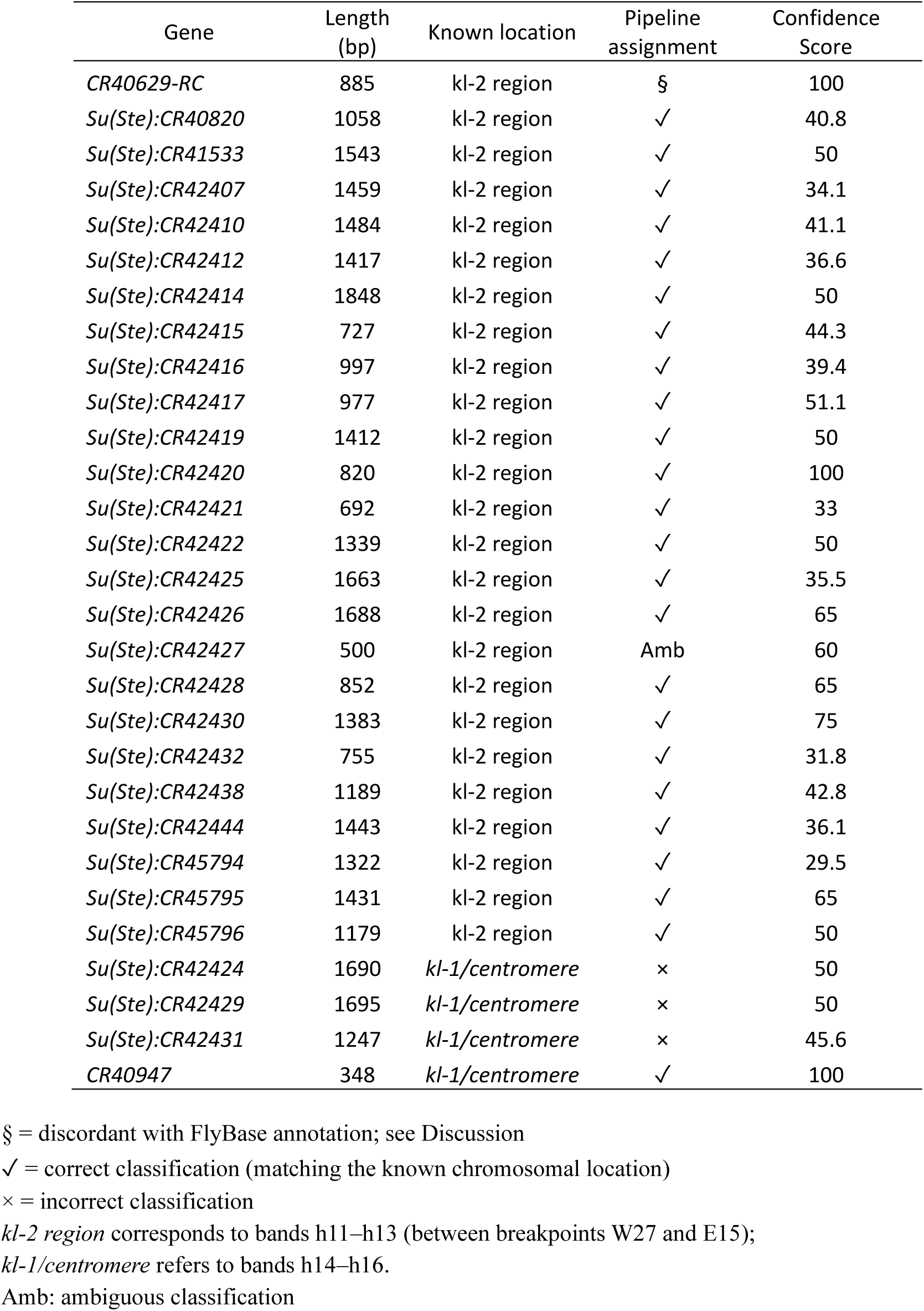
Results of the Digital Kennison classification for 29 Y-linked non-coding RNA sequences under optimized parameters. Pipeline assigned location, known location, and confidence score are shown.

| Gene | Length<br>(bp) | Known location | Pipeline<br>assignment | Confidence<br>Score |
| --- | --- | --- | --- | --- |
| *CR40629-RC* | 885 | kl-2 region | § | 100 |
| *Su(Ste):CR40820* | 1058 | kl-2 region | ✓ | 40.8 |
| *Su(Ste):CR41533* | 1543 | kl-2 region | ✓ | 50 |
| *Su(Ste):CR42407* | 1459 | kl-2 region | ✓ | 34.1 |
| *Su(Ste):CR42410* | 1484 | kl-2 region | ✓ | 41.1 |
| *Su(Ste):CR42412* | 1417 | kl-2 region | ✓ | 36.6 |
| *Su(Ste):CR42414* | 1848 | kl-2 region | ✓ | 50 |
| *Su(Ste):CR42415* | 727 | kl-2 region | ✓ | 44.3 |
| *Su(Ste):CR42416* | 997 | kl-2 region | ✓ | 39.4 |
| *Su(Ste):CR42417* | 977 | kl-2 region | ✓ | 51.1 |
| *Su(Ste):CR42419* | 1412 | kl-2 region | ✓ | 50 |
| *Su(Ste):CR42420* | 820 | kl-2 region | ✓ | 100 |
| *Su(Ste):CR42421* | 692 | kl-2 region | ✓ | 33 |
| *Su(Ste):CR42422* | 1339 | kl-2 region | ✓ | 50 |
| *Su(Ste):CR42425* | 1663 | kl-2 region | ✓ | 35.5 |
| *Su(Ste):CR42426* | 1688 | kl-2 region | ✓ | 65 |
| *Su(Ste):CR42427* | 500 | kl-2 region | Amb | 60 |
| *Su(Ste):CR42428* | 852 | kl-2 region | ✓ | 65 |
| *Su(Ste):CR42430* | 1383 | kl-2 region | ✓ | 75 |
| *Su(Ste):CR42432* | 755 | kl-2 region | ✓ | 31.8 |
| *Su(Ste):CR42438* | 1189 | kl-2 region | ✓ | 42.8 |
| *Su(Ste):CR42444* | 1443 | kl-2 region | ✓ | 36.1 |
| *Su(Ste):CR45794* | 1322 | kl-2 region | ✓ | 29.5 |
| *Su(Ste):CR45795* | 1431 | kl-2 region | ✓ | 65 |
| *Su(Ste):CR45796* | 1179 | kl-2 region | ✓ | 50 |
| *Su(Ste):CR42424* | 1690 | *kl-1/centromere* | × | 50 |
| *Su(Ste):CR42429* | 1695 | *kl-1/centromere* | × | 50 |
| *Su(Ste):CR42431* | 1247 | *kl-1/centromere* | × | 45.6 |
| *CR40947* | 348 | *kl-1/centromere* | ✓ | 100 |

§ = discordant with FlyBase annotation; see Discussion
✓ = correct classification (matching the known chromosomal location)
× = incorrect classification
*kl-2 region* corresponds to bands h11–h13 (between breakpoints W27 and E15);
*kl-1/centromere* refers to bands h14–h16.
Amb: ambiguous classification

Confidence scores reflect underlying classification difficulty. Single-copy genes and *Mst77Y* copies typically score high, reflecting the clean binary presence/absence patterns they produce across the deficiency panel, whereas repetitive *Su(Ste)* sequences score lower even when correctly classified. Of the five cases that disagreed with current FlyBase annotations, four (*CR42427*, *CR42424*, *CR42429*, and *CR42431*) received scores below 55, consistent with the score acting as a soft indicator of uncertainty rather than a strict exclusion threshold (*Supplementary Table S16* and *Supplementary Figure S5* in *Supplementary File S1*).

One exception is *CR40629-RC,* which received the maximum score (100) despite conflicting with its FlyBase annotation. This discrepancy persisted across all parameter settings, suggesting that the issue lies with the FlyBase annotation rather than with the pipeline classification. We examine this case below.

### Validation using known single-copy Y-linked genes

Single-copy genes provide the simplest benchmark for validation. We tested Digital Kennison on 13 well-characterized protein-coding genes spanning all six fertility regions, each originally assigned by wet-lab experiments using Kennison’s X-Y deficiency panel (*Supplementary Table S6 in Supplementary File S1*). All 13 were correctly assigned to their known cytological regions (Table 2; *Supplementary Table S10 in Supplementary File S1*). Ten received confidence scores ≥97.6, reflecting the clean binary presence/absence patterns expected for single-copy loci. Among them was *CG41561*, a Y-linked gene lacking any prior cytological assignment (Mahajan & Bachtrog, 2017), which was confidently mapped to the kl-1 region (confidence score 100). The remaining three cases illustrate more challenging scenarios: a gene with a close autosomal paralog (*FDY* /*vig2*) and two boundary-spanning genes (*PRY* and *Ppr-Y*).

*PRY* and *Ppr-Y* physically extend across adjacent fertility regions (Carvalho et al., 2000; Carvalho et al., 2001). Consequently, deletion of either region removes only part of the gene, producing intermediate occupancy in both corresponding databases rather than the near-zero signal expected for single-region loci. Digital Kennison recognizes this signature by identifying adjacent region pairs whose combined occupancies approximate full coverage. Accordingly, *PRY* showed intermediate occupancy in the kl-5⁻ (34%) and kl-3⁻ (66.2%) regions, whereas *Ppr-Y* showed the same pattern in the kl-2⁻ (53.5%) and kl-1⁻ (39.2%) regions. Both were correctly classified as boundary-spanning genes, with lower confidence scores (74 and 81.3) reflecting the absence of a fully binary occupancy profile.

*FDY* represents a different case: a single-copy gene with a near-identical autosomal paralog. Because *FDY* shares ∼98% nucleotide identity with *vig2*, BLAST-based analysis produces spurious cross-mapping, inflating occupancy across all databases. This issue was compounded by the failure of the kl-1⁻ assembly to recover the authentic *FDY* contig, generating an apparent absence signal that could not be distinguished from a genuine deletion using assembly data alone. The read-based branch resolved the case: mapping prefiltered Illumina reads (autosomal- and X-aligned reads are always removed, as described in Methods) directly to the *FDY* produced high coverage across all six fertility databases (86.6–100%), low coverage in V8 (21.3%), and none in V24. This pattern is consistent with a centromeric location and in agreement with the known position of *FDY* (Carvalho et al., 2015).

### The multi-copy *Mst77Y* gene cluster

We next tested the pipeline on the *Mst77Y* gene family, a tandemly arrayed cluster of 18 Y-linked copies located in the centromeric region, comprising 8 functional copies and 10 pseudogenes (the latter denoted as ψ). Copies vary in sequence divergence and structural completeness, including internal deletions (Krsticevic et al., 2010; Krsticevic et al., 2015). This dataset probes the pipelinés performance on multi-copy sequences.

All 18 copies were correctly assigned to the centromeric region (Table 2; *Supplementary Table S11 in Supplementary File S1*), with confidence scores ranging from 39 to 99.3, and were strongly influenced by gene completeness. Copies without major deletions were resolved by the standard centromeric pattern — high occupancy across all six fertility databases and low occupancy in V8 and V24 — whereas a subset of incomplete pseudogenes exhibited fragmented occupancy patterns and required relaxed or fragmentation-aware classification paths (*Supplementary Table S12 in Supplementary File S1*). The read-based branch independently classified all 18 copies, including pseudogenes, through the standard centromeric pattern (*Supplementary Dataset 2, sheets S8–S14*).

### Y-linked non-coding RNAs

We next assessed Digital Kennison on 29 Y-linked non-coding RNAs: 24 canonical *Su(Ste)* family members, three FlyBase-annotated *Su(Ste)* sequences previously mapped to the kl-1/centromere region (*CR42424, CR42429, CR42431*; included as test cases for minority paralog resolution), one *PCKR* (*pseudo-CK2β repeat*) family member (*CR40947*), and the transcript *CR40629-RC* (Table 3). Together, these comprise the most difficult test case in our benchmark, owing to their multi-copy organization, abundance asymmetry between paralogous clusters, and extensive sequence similarity across families. *Performance on the Su(Ste) repeat family*

*Su(Ste)* and *PCKRs* are closely related repeat families derived from the autosomal gene *Ssl* (*CK2β*). The canonical *Su(Ste)* family forms a large tandem cluster of hundreds of near-identical copies in the kl-2 region (h11–h13), whereas *PCKR*s form a smaller paralogous cluster located in the kl-1/centromere region (h14–h16; Chang & Larracuente, 2019). Because the two families retain substantial sequence homology, reads and BLAST alignments originating from one cluster can cross-map to the other, diluting the region-specific occupancy patterns expected from the translocation-deficiency panel.

Under optimized parameters, 24 of 29 sequences were correctly classified, one remained ambiguous, and four were misclassified (Table 3; *Supplementary Tables S13–S14 in Supplementary File S1*). Within the canonical *Su(Ste)* cluster, 23 of 24 were correctly assigned to kl-2, with only *CR42427* remaining unresolved. The *PCKR* sequence *CR40947* was correctly localized to the kl-1/centromere region. Three of the four misclassifications involved FlyBase-annotated *Su(Ste)* transcripts (*CR42424*, *CR42429*, and *CR42431*), which lie in the kl-1/centromere region (∼150 kb distal to *WDY*; *Supplementary Table S8 in Supplementary File S1*) but were assigned to kl-2. We return to this discrepancy in the Discussion. The fourth case, *CR40629-RC*, is treated separately below.

### CR40629-RC: evidence for an annotation error in FlyBase

*CR40629-RC* is currently annotated in FlyBase as a transcript located in the kl-2 region, yet Digital Kennison consistently assigned it to the kl-5 region with maximal confidence (score 100) across all parameter combinations tested (*Supplementary Table S12 in Supplementary File S1* and *Supplementary Dataset 2, sheets S15-S22*). To investigate this discrepancy, we extracted all R6 sequences >400 bp long and ≥80% identical to CR40629-RC from the R6 assembly (56 sequences total) and classified them using the optimized parameter configuration (*Supplementary Results, S4.1.5 in Supplementary File S1*).

The resulting distribution was strongly skewed toward kl-2, with *CR40629-RC* as the sole outlier: 52 sequences (92%) were assigned to kl-2, two to ks-2 with low confidence (28.1), and one yielded an unresolved centromere/kl-2 classification. *CR40629-RC* itself was the only sequence in the family assigned to kl-5, with a confidence score of 100. This separation indicates that the pipeline distinguishes *CR40629-RC* from its paralogs by differential occupancy across the deficiency panel, rather than by sequence similarity alone. A full account of the cross-assembly BLAST analysis supporting this finding is provided in the *Supplementary File S1* (*Supplementary Table S15*) and in the *Supplementary Dataset 2* (*sheet S23*).

Taken together, these findings indicate that *CR40629-RC* is structurally distinct from its closest paralogs and most likely resides in the kl-5 region rather than in the kl-2 region.

### Centromere contig classification

We applied Digital Kennison to the five *D. melanogaster* centromere contigs identified by Chang et al. (2019), one from each of the major chromosomes. The pipeline correctly identified the Y centromere contig (Y_Contig26, CenY) as centromeric, while the four autosomal centromere contigs returned as *Undetermined*, consistent with their non-Y origin.

Accurate classification of Y_Contig26 required reducing the hit-weight factor from its default value (2.0) to 0. Under default parameters, the high density of repetitive BLAST hits forced the identity threshold to 100%, suppressing occupancy below the classification threshold, preventing assignment despite a qualitatively correct centromeric pattern (*Supplementary Dataset S2*, *sheets S28–S29*).

Notably, reducing the hit-weight factor also caused one autosomal centromere contig (tig00057289, Cen2) to receive a spurious centromeric call from the BLAST branch. However, the confidence score (19.2) fell below the minimum threshold for single-branch classifications (50), and the lack of supporting read-based evidence meant the call was rejected, thereby preserving the correct *Undetermined* classification.

### Localization of Y-linked scaffolds across genome assemblies

#### Cross-assembly evidence for revised chromosomal assignment of R6 minor scaffolds

We next applied Digital Kennison to the minor scaffolds of the Release 6 (R6) assembly, many of which remain unlocalized or are assigned only to broad chromosomal regions. Because this set includes both Y-linked and non-Y sequences, we first screened all 1,870 scaffolds using the YGS method and retained the 904 with strong evidence of Y-linkage (P_VSC_UK ≥ 80%; *Supplementary Figure S2 in Supplementary File S1*). These candidate Y-linked scaffolds were then analyzed with Digital Kennison to assign them to specific fertility regions.

Of the 904 scaffolds, 674 (74.6%) received a regional assignment, with kl-1 accounting for the largest fraction (25.3%), followed by the centromere (15.2%), and kl-2 (12.7%); the remaining regions each accounted for 3.7–8.3% of assignments. One scaffold was classified as ambiguous (showing support for two adjacent regions), and 229 (25.3%) remained undetermined because their occupancy patterns could not be unambiguously assigned to any fertility region

The YGS pre-filtering step also revealed five scaffolds whose strong Y-linkage signal was inconsistent with their current chromosomal annotations. Although all are labeled as centromere-proximal scaffolds from chromosomes 2 or 3, each exhibits a strong Y-linkage signal (P_VSC_UK ≥ 99.9%) that is consistently recovered across four assemblies (R6, CL, ONT, LILAP). Three were independently annotated as Y-linked in the CL assembly, while the remaining two mapped to CL contigs that also showed strong Y-linkage (P_VSC_UK ≥ 95.4% in all four assemblies; *Supplementary Table S18 in Supplementary File S1* and *Supplementary Dataset 2, sheet S24*). Digital Kennison further assigned three of these five scaffolds to specific fertility regions (kl-3, kl-2, and ks-2; *Supplementary Table S18 in Supplementary File S1*).

All five scaffolds were unlocalized in the Release 5 assembly and acquired their current 2Cen/3Cen annotations only at the R6 release, suggesting that these assignments reflect placement decisions made during R6 construction rather than independent cytological or genetic evidence. We therefore flag these five scaffolds as candidates for reassignment to the Y chromosome. A systematic re-evaluation lies beyond the scope of this study; further detail, including per-assembly P_VSC_UK values, is provided in the Supplementary Results (*Supplementary Table S18 in Supplementary File S1; Supplementary Dataset 2, sheet S24*).

#### Classification of unlocalized Y-linked scaffolds in two de novo assemblies

We tested Digital Kennison with two de novo assemblies lacking any scaffolding based on prior information (ONT and PacBio HiFi; *Supplementary Methods, S3.5.4 in Supplementary File S1*). In each case, scaffolds with strong Y-linkage signal (P_VSC_UK ≥ 80%) were identified using the YGS method. The pipeline assigned fertility regions to 37 of 61 candidates (60.7%) in the PacBio HiFi assembly and to 17 of 50 (34%) in the ONT assembly. In both datasets, most assignments required accepting the read-based classification alone, as BLAST occupancy was too low or too uniform to produce a regional call; without this allowance, the proportion of localized scaffolds dropped to 26.1% and 16%, respectively (*Supplementary Table S20 in Supplementary File S1*).

Although the HiFi assembly yielded a substantially higher proportion of classified scaffolds than the ONT assembly (60.7% vs. 34.0%), the difference was much smaller when measured by sequence length (65.5% vs. 54.7%). Both assemblies contributed nearly the same amount of classified sequence (∼10.5 Mb; *Supplementary Table S25 in Supplementary File S1*). In the ONT assembly, much of the unclassified sequence is concentrated in a few very large scaffolds, including a single 3.89 Mb scaffold (ptg000015l), whereas in the HiFi assembly the unclassified sequence is distributed among a larger number of smaller scaffolds (*Supplementary Dataset 2, sheets S31 and S34*).

The lower ONT classification rate was not explained by repeat content, scaffold length, read-mapping identity (*Supplementary Tables S21–S23 in Supplementary File S1*), or assembly chimerism. Windowed composition analysis identified only a small number of chimeric scaffolds in both assemblies, at similar frequencies (*Supplementary Figure S4 in Supplementary File S1*), ruling out chimerism as the cause of the observed difference.

### Application to recently transferred Y-linked sequences

Tobler et al. (2017) identified 21 recent Y-linked sequence transfers in *D. melanogaster* by short-read mapping. Although Y-linkage was established for these transfers, their precise location within the Y chromosome remained unknown, except for *FDY*, previously mapped to the centromeric region (Carvalho et al., 2015). After collapsing co-transferred sequences and excluding two that failed the alignment chaining step (*Supplementary Methods, section S3.5.3 in Supplementary File S1*), we applied Digital Kennison to 16 transfers to assign them to specific Y-chromosome fertility regions, using the known *FDY* localization as an internal positive control.

The pipeline localized 7 transfers to specific fertility regions, four of them with high-confidence scores, and three within the previously defined reliable classification range (55-80) (Table 4; *Supplementary Table S16*). As an internal positive control, the *FDY* region (Dmel_3R_20.95) showed complete concordance across its four co-transferred genes (*vig2, Bili, Clbn*, and *Mocs2–CG42503*), which converged on a single ∼29 kb interval (Y:3,021,897–3,051,262; *Supplementary Table S20 in Supplementary File S1*). The pipeline recovered the expected centromeric assignment for *FDY* with an aggregate confidence score of 73.3 (*Supplementary Dataset 2, sheet S27*).

**Table 4.** Digital Kennison classification of consensus gene transfers to the Y chromosome reported by Tobler et al. (2017). Categories were assigned by aggregating Y-linked hits per donor region: *Resolved* (single dominant location with high support), *Multicopy single region* (multiple hits with consistent dominant location), *Dispersed* (hits across multiple regions, no dominant assignment), *Insufficient evidence* (no reliable classification). Donor coordinates are FlyBase R5; final modal indicates the dominant inferred Y location. See Methods and *Supplementary Methods* for category definitions and aggregation thresholds.

| category | donor | consensus | Final modal | Final mean confidence score |
| --- | --- | --- | --- | --- |
| Resolved | 2L:4456606-4457002 | Dmel_2L_4.46 | kl-1 | 95 |
|  | 2L:19936154-19939442 | Dmel_2L_19.94 | kl-2 | 85.1 |
|  | 3L:24503593-24524235 | Dmel_3L_24.3 | ks-1 | 55.8 |
|  | 3R:17042662-17042936 | Dmel_3R_17.04 | centromere | 92.3 |
|  | Dmel_3R_20.95 (FDY region) | Dmel_3R_20.95 | centromere | 73.3 |
|  | X:12656236-12657385 | Dmel_X_12.7 | centromere | 82.9 |
| Multicopy single region | 2R:2350137-2350570 | Dmel_2R_2.32 | kl-2 | 72.7 |
| Dispersed | 2L:22752196-22752705 | Dmel_2L_22.75 | multi-region | 70.5 |
|  | 2R:99735-100642 | Dmel_2R_0.09 | multi-region | 68.8 |
|  | 2R:566974-568706 | Dmel_2R_0.57 | multi-region | 61.1 |
|  | 3L:23412979-23413541 | Dmel_3L_23.41 | multi-region | 86 |
|  | 3L:24301186-24302152 | Dmel_3L_24.3 | multi-region | 71.6 |
|  | 3L:24506237-24507024 | Dmel_3L_24.3 | multi-region | 65 |
|  | X:21824921-21828314 | Dmel_X_21.83 | multi-region | 61.2 |
| Insufficient evidence | 2R:737727-739380 | Dmel_2R_0.57 | Undetermined | 75 |
|  | 3L:24432130-24442098 | Dmel_3L_24.3 | Undetermined | 41.4 |

multi-region: no single location predominates
Undetermined: when no assignment could be made

The remaining nine transfers could not be localized to a single region. These showed Y-linked hits dispersed across multiple fertility regions or lacked sufficient signal for classification, consistent with sequences present on the Y as short, scattered copies rather than a single contiguous insertion (Table 4; *Supplementary Dataset 2, sheets S26* and *S27*).

## Discussion

Despite remarkable advances in sequencing technologies, heterochromatic regions remain systematically underrepresented in *Drosophila* genomic assemblies. While single-molecule platforms like PacBio and Oxford Nanopore have enabled essentially complete assemblies of the euchromatic *Drosophila* genome, heterochromatin — including the entire Y chromosome — remains fragmented and incomplete. This limitation is unlikely to disappear soon because all current long-read technologies exhibit strong bias against simple satellites (Carvalho et al., 2026), which constitute a major fraction of *Drosophila* heterochromatin, and are particularly abundant in the Y chromosome (Lohe et al., 1993). As a result, genome assemblies alone are often insufficient to determine the location of newly identified Y-linked sequences. Classical Kennison mapping can provide this information, but its practical demands—maintaining translocation stocks, performing crosses, and conducting PCR assays—have limited its routine application. Consequently, several Y-linked loci remain unmapped, including the *CG41561* gene (Mahajan & Bachtrog, 2017). Digital Kennison bridges this gap by recasting the classical mapping as a computational analysis that requires only sequence data, not physical access to the Kennison strains.

The most immediate consequence of this shift is that fertility-region mapping becomes a routine computational task. In this study, *CG41561*, a Y-linked gene identified by Mahajan & Bachtrog (2017) but never cytologically mapped, was assigned to the kl-1 region with maximum confidence. Of 16 recent autosome-to-Y transfers described by Tobler et al. (2017), 7 were localized to specific fertility regions, including 4 with high confidence. Applied to the 904 Y-linked minor scaffolds of the Release 6 assembly, the pipeline assigned 674 (74.6%) to specific Y-chromosome regions, substantially expanding the fraction of Y-linked sequences with regional annotations. These results illustrate the pipeline’s primary intended use: rapid localization of sequences that would otherwise require extensive genetic analysis.

Routine computational mapping also makes it practical to re-examine existing annotations, something rarely done because classical mapping experiments are seldom repeated once an assignment has been made. Two findings from this study illustrate this. The first concerns *CR40629-RC*, which Digital Kennison consistently mapped to the kl-5 region despite its current FlyBase annotation placing it in the kl-2 region. This discrepancy, combined with the consistent kl-2 localization of its 55 closest paralogs, is more parsimoniously explained by an annotation error than by a pipeline misclassification. The second concerns five R6 minor scaffolds currently annotated as 2Cen or 3Cen. All five showed strong Y-linkage signals across four independent assemblies, and three were further localized to specific Y fertility regions by the pipeline. Because these scaffolds were unlocalized in Release 5 and acquired their current autosomal annotations only in Release 6, they represent plausible candidates for reassignment to the Y chromosome.

These examples also highlight an important principle: No mapping strategy is free of artifacts, particularly in highly repetitive genomic regions. The placement of *Pp1-Y1* illustrates this: A read-depth analysis by Chang & Larracuente (2019) suggested a location near the *Su(Ste)/PCKR* family, conflicting with earlier PCR-based mapping to the kl-5 region (Carvalho et al., 2001, 2015). Digital Kennison independently recovered the kl-5 assignment, adding a third line of evidence in its favor despite the unresolved discrepancy. The same logic underlies the reassignment of *CR40629-RC* and the candidate Y-linked scaffolds reported here: these conclusions rely not on the pipeline alone but on its concordance with independent evidence, including paralog distributions and YGS-supported Y-linkage across multiple assemblies.

The cases examined here also define the conditions under which the pipeline reaches its limits, although these limits are not specific to the computational approach. Closely related paralogs, for example, confound any method that relies on sequence similarity, including PCR, FISH, and read mapping, because the underlying ambiguity lies in the sequences themselves rather than in any particular detection strategy. The two paralog-confounded cases encountered in this study illustrate distinct failure modes and how the dual-evidence framework addresses each.

*FDY* illustrates a data loss problem. Its nearly identical autosomal paralog, *vig2*, inflates BLAST occupancy across all databases, while failure to assemble the authentic FDY copy in the kl-1^-^ strain produces an artifactual absence pattern that BLAST alone cannot distinguish from a genuine deletion. The read-based branch resolved the ambiguity by operating on sequencing reads rather than assembled contigs, thereby bypassing both paralog inflation and assembly artifacts. The *Su(Ste)* transcripts illustrate a different problem: signal dominance. Because the canonical *Su(Ste)* family is highly concentrated in the kl-2 region, BLAST alignments to these abundant paralogs dominated the occupancy profile, biasing assignments toward kl-2 regardless of the query’s true position. The persistent inability to fully resolve the three kl-1/centromere *Su(Ste)* sequences (*CR42424, CR42429, CR42431*) under any tested configuration demonstrates the broader limitation posed by abundance-asymmetric paralog families for sequence-based mapping inference. Importantly, the resulting misclassifications were directional rather than random: their tendency to shift from kl-2 toward the centromere is consistent with secondary signal from the pericentromeric *PCKR* cluster being recovered when parameter settings cannot fully suppress cross-mapping.

Highly repetitive and centromeric sequences present a related challenge. Correct classification of the Y centromere (Y_Contig26) required relaxing the hit-weight penalty to compensate for the hundreds of repetitive BLAST hits produced by shared centromeric retroelements. Under those same permissive settings, one autosomal centromere contig (tig00057289) also generated a spurious Y-linked signal, but its low confidence score and lack of read-based support prevented an incorrect final classification. This outcome reflects a biological constraint: all five *D. melanogaster* centromeres share a common complement of retroelements — most notably *G2/Jockey-3*, the only element present on all five and indistinguishable, phylogenetically, from genome-wide copies (Chang et al. 2019). Sequence-based localization of the Y centromere is therefore bounded by inter-centromere sequence sharing.

Input data quality also determines classification success in de novo assemblies, although the magnitude of this effect depends on how resolution is measured. By scaffold count, the pipeline localized 60.7% of candidate Y-linked scaffolds in the HiFi assembly, but only 34.0% in the ONT assembly. By assembled sequence, however, the difference narrowed to 65.5% versus 54.7%, with nearly identical amounts of sequence classified (∼10.5 Mb in each assembly; Table S25). Most of the discrepancy between these metrics is explained by a small number of very large Undetermined scaffolds in the ONT assembly, indicating that the difference between assemblies reflects, in large part, how sequence is partitioned among contigs rather than the amount of Y sequence the pipeline is able to resolve. The remaining gap is not explained by read quality, repeat content, scaffold length, or scaffold chimerism (*Supplementary Tables S21–S24 and Supplementary Figure S4 in Supplementary File S1*), and its cause remains unresolved.

In each of these cases (centromeric cross-mapping, chimeric scaffolds, paralog dominance), the practical limit is imposed by the input data, and the confidence score conveys that limit to the user. Two general limitations remain. First, classification depends on the deficiency-panel assemblies, and a sequence that is present in a strain but fails to assemble produces false absence patterns. The read-based branch substantially mitigates this problem but cannot eliminate it entirely. Second, the method is designed to localize sequences represented by a dominant contiguous Y-linked locus and therefore cannot reliably resolve cases in which Y representation consists primarily of numerous short fragments dispersed across multiple fertility regions, as observed for 9 of the 16 Y-linked sequence transfers analyzed here (Tobler et al., 2017).

The Kennison translocation lines have played a central role in shaping our understanding of the *Drosophila* Y chromosome. Beyond their use as mapping tools, they enabled the generation of males deficient for specific Y regions and thereby provided direct access to the phenotypic consequences of fertility-factor losses. A direct line connects these strains to the identification of *kl-3* and *kl-5* as axonemal dynein heavy-chain loci (Hardy et al., 1981; Goldstein et al., 1982) and to the first molecular characterization of a Y-linked protein-coding gene in *Drosophila* (Gepner & Hays, 1993), which laid the foundation for localizing the remaining Y-linked genes (Carvalho et al., 2000, 2001, 2015; Mahajan & Bachtrog, 2017). These strains remain relevant today: a complete telomere-to-telomere assembly of the *Drosophila* Y chromosome is still an unsolved challenge (Carvalho et al., 2026), and the pipeline presented here extends the utility of the Kennison lines by removing the practical barriers that have limited their routine use. In doing so, it provides a rapid and scalable framework for localizing newly discovered Y-linked genes, transcripts, and repetitive sequences while preserving the power of classical translocation mapping

## Data and Code Availability

*Supplementary File S1* (methods, results, figures, and tables) accompanies the manuscript through the journal’s submission system. Two large supplementary datasets (*Supplementary Datasets 1 and 2*) are archived at Zenodo due to file size: https://doi.org/10.5281/zenodo.21367000. *Supplementary Dataset 1* contains the YGS output tables for all translocation-strain assemblies (*sheets S1–S7*) and per-scaffold PVSC_UK values for the five assemblies used to validate R6 minor-scaffold Y-linkage (*sheets S8–S12*). *Supplementary Dataset 2* contains the cross-assembly BLAST tables and centromere-contig analyses supporting the main-text findings.

The Digital Kennison pipeline is available at https://github.com/fabianauno/digital_kennison and archived at Zenodo (https://doi.org/10.5281/zenodo.21271417). The associated data are archived at Zenodo as three related records:

- Kennison deficiency-panel BLAST databases: https://doi.org/10.5281/zenodo.21280260
- Prefiltered short reads from the Kennison X-Y translocation panel: https://doi.org/10.5281/zenodo.21284273
- Y -linked scaffold identifier lists (gi_list files) derived from YGS analysis: https://doi.org/10.5281/zenodo.21296560

Raw whole-genome sequencing reads for the eight Kennison X-Y translocation strains and for the Drosophila melanogaster ISO1 reference female are archived at NCBI SRA under BioProject PRJNA1493028. All Zenodo records are grouped in the Digital Kennison community: https://zenodo.org/communities/digital-kennison.

## Acknowledgments

We thank Felipe Bastos Rocha, Thyago Vanderlinde, Leonardo Barbosa Koerich, and members of our laboratory for valuable discussions and suggestions throughout this study.

